# AutoLead: An LLM-Guided Bayesian Optimization Framework for Multi-Objective Lead Optimization

**DOI:** 10.1101/2025.08.19.671029

**Authors:** Yiming Zhang, Jun Jin Choong, Kaushalya Madhawa, Keisuke Ozawa

**Affiliations:** Department of Computational Biology and Medical Sciences, Graduate School of Frontier Sciences, The University of Tokyo, Japan; SB Intuitions, Japan

**Keywords:** Drug discovery, Lead optimization, Bioinformatics, Large Language Models, Bayesian optimization

## Abstract

The process of lead optimization in drug discovery is a complex, multi-objective challenge that remains a major bottleneck in the development of new therapeutics. Traditional approaches often struggle to efficiently explore the vast chemical space while simultaneously optimizing multiple, and sometimes conflicting, molecular properties. In this work, we present AutoLead, a novel framework that integrates Large Language Models (LLMs) with multi-objective Bayesian optimization to tackle this challenge. By leveraging the chemical reasoning capabilities of LLMs, AutoLead effectively guides the search for novel drug-like molecules that satisfy multiple objectives. We evaluate our approach on two molecule optimization tasks, achieving state-of-the-art results. Furthermore, we introduce a new benchmark dataset designed around a more realistic lead optimization scenario, where the task is to modify compounds that violate Lipinski’s Rule of Five to simultaneously meet all criteria and improve their QED score. Through extensive experiments and a detailed case study, we demonstrate the potential of combining LLMs with black-box optimization techniques for more efficient and practical drug discovery.

## Introduction

Lead optimization is a critical stage in drug discovery, where chemists iteratively modify hit compounds to improve their pharmacokinetic properties while ensuring efficacy and safety (1, 2). Unlike early-stage virtual screening, lead optimization involves complex, multi-objective trade-offs, such as simultaneously improving solubility, permeability, and drug-likeness, under strict medicinal chemistry constraints (3, 4). Traditional approaches often rely on expert heuristics or rule-based systems to make localized edits (5), which typically operate in oversimplified settings and fail to scale in the face of combinatorial chemical space and tightly coupled objectives (3, 6).

Recent advancements in large language models (LLMs) have opened promising avenues for pipelining molecular optimization (6–9). This has led to two main strategies. One involves developing domain-specific LLMs that are pre-trained on vast scientific corpora to enhance their chemical reasoning (10–12). Specifically for molecular optimization, DrugAssist (13) achieved leading results by instruction-tuning an LLM on a large dataset for multi-property optimization. These works highlight the potential of LLM-guided optimization but often lean on specialized models or extensive fine-tuning.

While these approaches highlight the power of LLMs in cap-turing chemical reasoning and facilitating property-guided edits, they also exhibit limitations, particularly their reliance on large-scale instruction-finetuning datasets tailored for molecule editing. The other adapts powerful general-purpose models for chemical tasks. ChatDrug (14) introduced a conversational framework to iteratively edit molecules through prompt design and dialogue. However, ChatDrug relies on an external molecular database for retrieval, a dependency that inherently restricts its exploration to known chemical space and limits its ability to generate potential novel candidates (15). Moreover, the tasks and datasets used there primarily focus on single-objective optimization or loosely coupled dual-property scenarios, falling short of the complex, tightly constrained multi-objective requirements encountered in realistic lead optimization workflows.

To overcome these limitations, we introduce a new framework—**AutoLead**—that combines the strengths of LLMs with the principled decision-making capabilities of Bayesian Optimization (BO), without requiring any external retrieval database or domain-specific training data. In our approach, the LLM first proposes a diverse set of candidate molecules by leveraging its inherent knowledge of chemical structure and function. These initial proposals serve as seeds for BO, which then guides the optimization process by selectively exploring or exploiting candidates based on historical performance and uncertainty estimates modeled by Gaussian Processes (GPs) (16, 17). This integration allows AutoLead to dynamically alternate between LLM-driven exploration and GP-guided exploitation, enabling efficient navigation of the chemical space under multi-objective constraints.

In practical drug discovery, lead optimization often begins with modifying hit compounds that violate key drug-like properties, such as Lipinski’s Rule of Five (18, 19), to simultaneously satisfy multiple pharmacokinetic constraints and improve drug-likeness metrics like QED. To evaluate our method in this realistic setting, we compile a new benchmark dataset derived from the HiQBind (20) dataset, explicitly designed for lead optimization where the objective is to transform molecules that fail one or more Lipinski criteria into candidates that satisfy all rules while enhancing their QED scores.

Our key contributions are summarized as follows:

- We propose AutoLead that integrates general-purpose LLMs with multi-objective BO specifically designed to generate and refine molecules under realistic pharmacokinetic constraints, without target-specific tuning or retrieval databases.
- We demonstrate that AutoLead achieves state-of-the-art performance on three benchmarks, ChatDrug-200, DrugAssist-500, and our newly introduced LipinskiFix-1000, highlighting its broad applicability and effectiveness in complex multi-objective settings.
- We construct LipinskiFix-1000, a challenging benchmark dataset derived from HiQBind (20), designed to evaluate lead optimization methods on tasks that require repairing non-compliant molecules while improving overall drug-likeness.

Together, these contributions showcase how combining LLMs’ generative flexibility with BO’s principled exploration can yield a powerful and data-efficient solution to real-world lead optimization.

## Materials and methods

### A. Overview

We tackle molecular lead optimization, where given an initial molecule, the goal is to generate a sequence of improved molecules by optimizing multiple drug-relevant properties such as drug-likeness (QED), lipophilicity (logP), and Lipinski compliance, addressing the challenge of multi-objective optimization that existing methods often struggle with.

AutoLead addresses this problem via a hybrid *LLM-in-the-loop Bayesian optimization* framework (Figure 1) that couples the generative chemical reasoning capabilities of general-purpose LLMs with uncertainty-aware optimization using a GP surrogate model. We also provide a mapping strategy between SMILES strings and feature vectors to mitigate the difficulty of incorporating BO into our hybrid scheme.

**Fig. 1.**
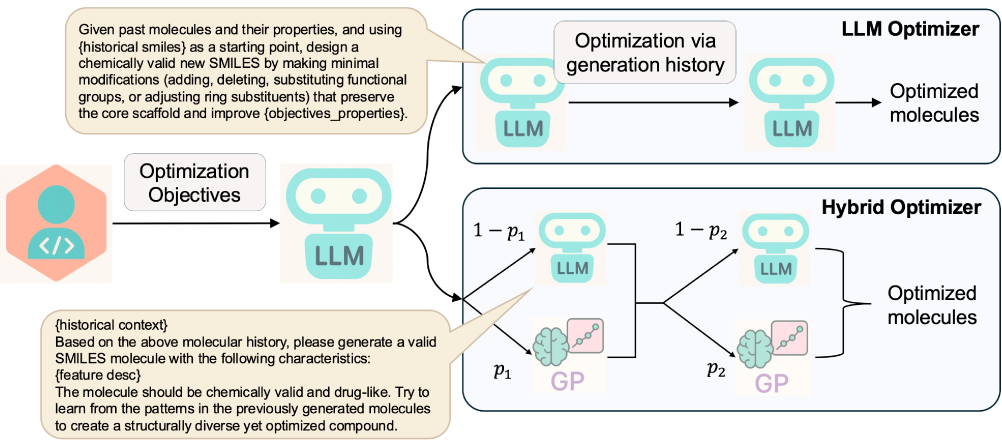
Overview of the AutoLead framework. The system combines an LLM-based optimizer that leverages historical molecular properties and a hybrid optimizer integrating both LLM and GP components. Warmstarting uses the LLM to propose initial molecules with standard scaffolds. Subsequent optimization balances LLM chemical reasoning with uncertainty-aware BO to progressively refine molecules toward multi-objective drug-like criteria.

### B. Problem Formulation

Let 𝒮 denote the set of valid SMILES (21) strings and let

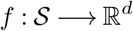

be a property evaluator that returns a vector of target properties such as QED, logP, tPSA, number of hydrogen bond donors/acceptors and molecular weight. We aim to solve the optimization problem:

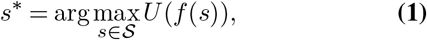

where *U* (·) aggregates the *d* objectives, and is by default taken as a uniform average: 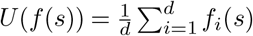.

### C. Surrogate Objective and LLM-driven Inverse Mapping

Since BO cannot directly optimize over discrete SMILES strings, we introduce a continuous descriptor space:

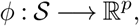

where *ϕ*(*s*) maps a molecule *s* to a *p*-dimensional vector of normalized RDKit (22) descriptors. To ensure all values fall within the range [0, 1] for GP-based Bayesian Optimization, each descriptor is divided by a fixed normalization constant (shown in parentheses). Specifically, we use 10 molecular properties: molecular weight (500 Da), logP (5.0), polar surface area (200 Å^2^), number of hydrogen bond donors (10), number of hydrogen bond acceptors (10), number of rotatable bonds (20), ring count (10), number of aromatic rings (5), number of saturated rings (5), and sp^3^ carbon fraction (which is inherently bounded within [0, 1]. This normalization mitigates heterogeneous units and improves GP conditioning and acquisition optimization, in line with prior recommendations that objective scales and trust-region potentials are dynamic during optimization (23)).

This 10-dimensional vector provides a pharmacologically meaningful and normalized space for latent space optimization. It is important to note that *f* is a property evaluator (e.g., computing logP), whereas *ϕ* is used to represent molecules for optimization.

A technical challenge, however, is converting an optimized point from this latent space back into a valid chemical structure (a SMILES string). This inverse mapping, *ϕ*^−1^, is generally intractable. To overcome this, we propose a training-free LLM-driven decoder, denoted as *ϕ*^−1^. Instead of attempting a direct conversion from latent space, our approach leverages the generative power of an LLM (24). We translate the optimized latent vector *x* ∈ [0, 1]^*p*^(*p* = 10) into a human-readable text prompt. For instance, the vector is verbalized into a sentence such as, “Generate a molecule with a molecular weight of approximately 310 Da, a logP of 3.5, …,” instructing the LLM to produce a molecule that satisfies these 10 target properties. The optimization process is therefore approximately realized by:

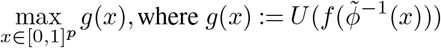

In this framework, BO efficiently searches for the optimal vector in the continuous latent space, and our LLM-driven decoder acts as a bridge, translating these optimized vectors back into tangible molecules. Finally, the generated molecules are evaluated by the property evaluator f to verify if they meet the desired optimization goals.

### D. Bayesian Surrogate Modeling

As in (25), given observations up to step (*t*−1), we define the data points as:

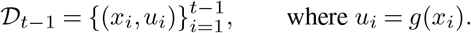

We place a GP prior over the utility function *U* (*x*) and obtain the following posterior distribution:

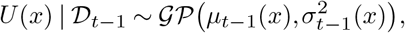

where the predictive mean and variance are given by:

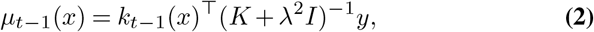

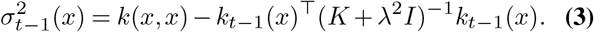

Here, *K* ∈ ℝ^(*t*−1)*×*(*t*−1)^ is the Gram matrix of the training inputs, 𝒟_*t* − 1_, where each entry is defined as *K*_*ij*_ = *k*(*x*_*i*_, *x*_*j*_) for *i, j* ∈ {1,…, *t* − 1} . The vector *k*_*t* − 1_(*x*) = [*k*(*x, x*_1_), …, *k*(*x, x*_*t* − 1_)]^⊤^ contains the covariances between the test point *x* and the training inputs. The vector *y* = [*u*_1_, …, *u*_*t* − 1_]^⊤^ stores the observed utility values. We use a radial basis function kernel *k*(·,·) as the covariance function.

### E. LLM-in-the-Loop Optimization Strategy

We employ a transient hybrid strategy (25) to gradually transition from LLM-based exploration to BO-based exploitation. The probability of using BO at round *t* is:

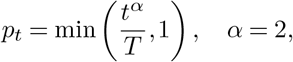

where *T* is the total iteration budget. Let *z*_*t*_ ∼ Bernoulli(*p*_*t*_). If *z*_*t*_ = 1 and |𝒟_*t*−1_ | ≥ *τ*, a threshold size of data points for switching the strategy, we invoke BO using our LLM decoding; otherwise, the LLM generates the molecule:

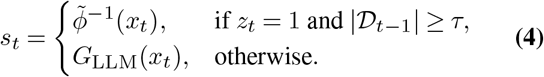

Here, 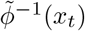 denotes molecule generation via the BO proposal obtained by maximizing the acquisition function in descriptor space, while *G*_LLM_ denotes a direct molecule generation by the LLM. We set *τ* = 3 so that BO starts only after at least three data points have been collected.

#### Acquisition Function

The BO step maximizes the upper confidence bound (UCB) acquisition function:

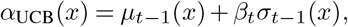

where

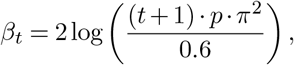

and *p* denotes the feature dimension, as also implemented in (25).

### F. Multi-Objective Handling

1. **Weighted Utility:** 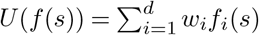, with default *w*_*i*_ = 1*/d*.
2. **Pareto Tracking:** maintain a Pareto front of non-dominated solutions (26).

### G. Overall Optimization Loop

Algorithm 1 outlines the overall optimization procedure of AutoLead. Starting from an initial molecule *s*_0_, the algorithm iteratively alternates between LLM-based exploration and BO-guided exploitation, controlled by a Bernoulli variable. At each step, a new candidate is generated, evaluated, and added to the historical optimization trajectory. The surrogate model and Pareto front are continuously updated, and the final output is a set of non-dominated molecules after *T* iterations.

#### Algorithm 1 AutoLead

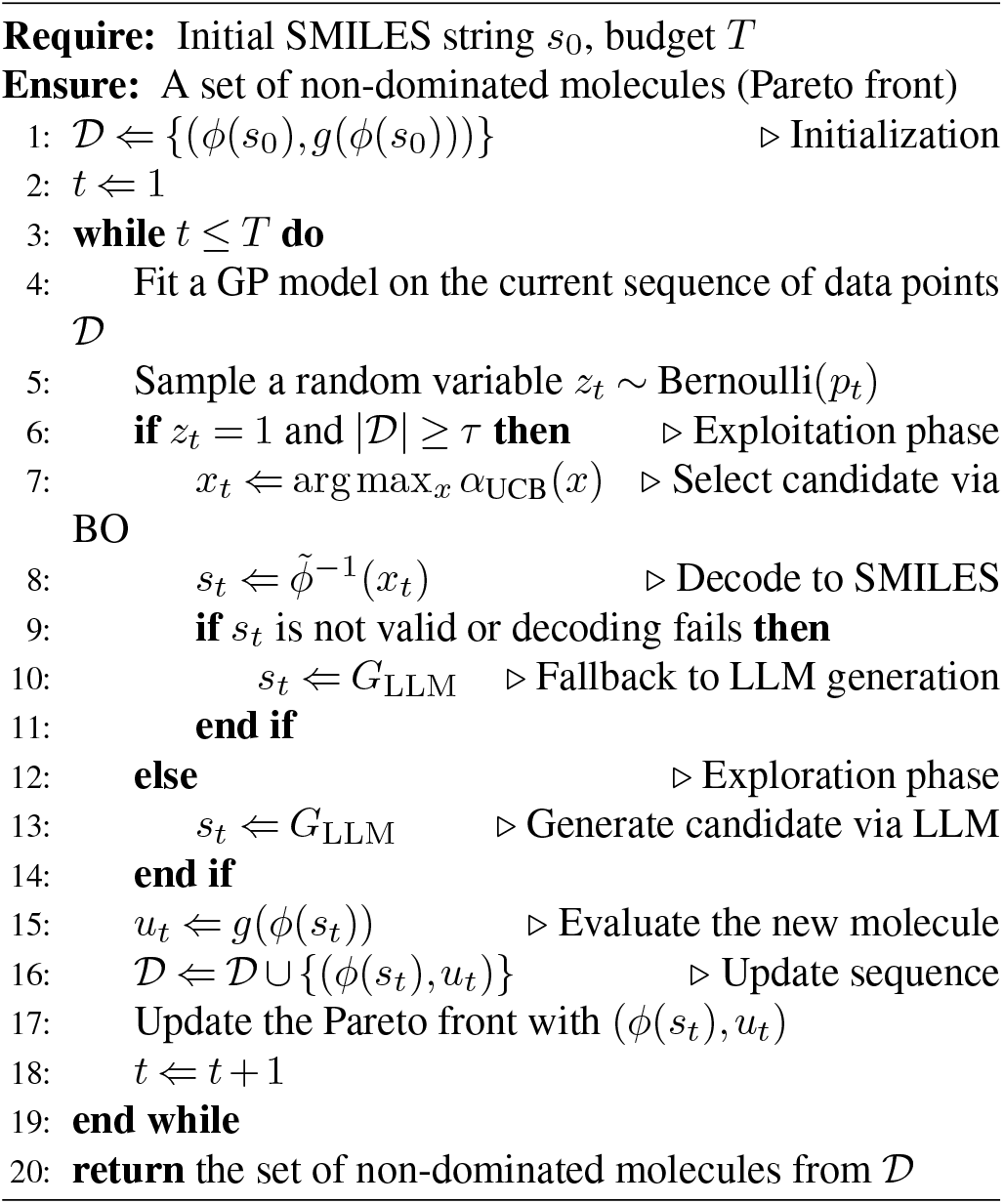

### H. Benchmark datasets

We evaluate AutoLead on multiple molecular optimization datasets covering diverse optimization scenarios, from single-objective property optimization to complex multi-objective lead optimization tasks.

#### H.1. LipinskiFix-1000

We further introduce LipinskiFix-1000, a benchmark dataset designed to evaluate molecular optimization strategies on realistic drug-like compounds. It consists of 1,000 unique ligands extracted from the high-quality HiQBind (20) database, each bound to experimentally validated protein targets. These molecules were selected for violating at least one of Lipinski’s Rule of Five criteria, reflecting common challenges in early drug discovery. To ensure meaningful optimization, we applied additional quality constraints on molecular weight, lipophilicity, and QED scores, while limiting structural complexity. Each task targets restoring Lipinski compliance as the primary objective and maximizing QED as a secondary goal, providing a rigorous testbed for multi-objective molecular design.

#### H.2. Dataset construction

We start with the high-quality HiQBind database, which provides experimentally validated protein–ligand complexes with curated binding affinities. We filter for ligands that:

- have complete property annotations (MW, LogP, HBD, HBA, QED),
- contain at most 50 heavy atoms to ensure manageable complexity,
- violate at least one Lipinski rule (MW *>* 500, LogP *>* 5, HBD *>* 5, or HBA *>* 10), and additionally satisfy quality checks (QED ≥ 0.1, MW ≤ 800, LogP ≤ 8) to maintain feasible optimization targets.

We remove duplicates based on SMILES strings and random sample 1,000 ligands to form LipinskiFix-1000. Each entry is paired with its protein context and annotated with detailed property values, violated rules, and explicit primary (Lipinski compliance) and secondary (QED maximization) optimization objectives.

#### H.3. Molecular Property Optimization Dataset

We evaluate AutoLead on two standard molecular property optimization benchmarks from ChatDrug and DrugAssist that give diverse optimization scenarios.

#### H.4. ChatDrug-200

We use 200 molecules curated from the ChatDrug framework, which systematically investigates conversational drug editing with LLMs. These molecules cover diverse single- and multi-objective optimization tasks, such as adjusting LogP, QED, permeability (mesured by TPSA), and modifying hydrogen bond acceptor/donor counts, under loose and strict improvement thresholds Δ.

#### H.5. DrugAssist-500

We additionally evaluate on 500 molecules curated from the DrugAssist benchmark, designed to test interactive multi-property optimization. Tasks include simultaneous optimization of properties like BBBP, hERG inhibition, solubility, and QED, often requiring target values to lie within specified ranges while preserving structural similarity. Since RDKit can not directly compute BBBP and hERG inhibition, we focus on solubility, QED, and acceptor/donor tasks for optimization under both loose and strict improvement threshold (Δ) constraints.

### I. Evaluation Metrics

We adopt the evaluation metrics used in (13, 14) to assess AutoLead’s performance across various optimization scenarios:

#### Success Rate

Percentage of molecules successfully optimized to meet specified thresholds.

#### Valid Rate

Percentage of chemically valid molecules.

## Results and Discussion

This section presents our experimental results. We begin by describing the experimental setup, including the datasets and baseline models. We then report and analyze the performance of our proposed method.

### Experimental Setup

To ensure the reliability of our results, all methods were evaluated over 3 independent runs for each task. We report the mean and standard deviation across these runs.

#### Specifications for AutoLead

We evaluate three variants of AutoLead to specifically demonstrate how the hybrid strategy improves performance:

- **LLMO (LLM Optimizer)**: An LLM-guided approach that leverages the large language model throughout all iterations to propose candidate molecules based on learned chemical priors.
- **BO (Bayesian Optimizer)**: A special case of the hybrid strategy where BO is used in every round (*z*_*t*_ ≡ 1), allowing the surrogate model to fully guide the optimization from start to finish.
- **HO (Hybrid Optimizer)**: A transient strategy that combines LLM exploration and Gaussian Process-based Bayesian Optimization (BO). It initially relies on the LLM to gather diverse molecules and switches to BO once a minimum sequence size (*τ* = 3 points) is reached, allowing exploitation of the surrogate model (25).

For both variants, we set the maximum number of optimization iterations *T* to 10 for each molecule. The process stops early if a molecule satisfying the target property threshold is found, or continues until the iteration limit is reached. All prompt templates used in this work are provided in the Supplementary Data.

### Baselines Methods

The baselines (27) are based on MegaMolBART (28), a pretrained auto-regressive model. Baselines include Random, PCA, High-Variance, GS-Mutate (15), and MoleculeSTM (27) with SMILES and ChatDrug with GPT-3.5 Turbo and GPT-4o (29) as the backbone LLMs. For the ChatDrug baselines, we set the number of conversation rounds to 10, and reran their implementation to be precisely aligned with the evaluation metrics, as also noted in (13). For the other baselines, we adopted the values reported in (14).

### Performance on the ChatDrug-200 dataset

Our proposed hybrid optimization method, AutoLead-HO, demonstrates a commanding performance on the ChatDrug-200 benchmark, significantly outperforming both existing state-of-the-art methods and its own ablation variants, AutoLead-LLMO and AutoLead-BO.

In the single-objective setting (Table 1), AutoLead-HO consistently achieves top success rates, especially under the most stringent test conditions. For instance, on challenging tasks such as increasing the hydrogen bond acceptor count (HBA+) or optimizing for QED+, our method significantly outperforms all baselines, including the strong ChatDrug with GPT-4o model. This highlights the robustness of our hybrid strategy in focused optimization scenarios.

**Table 1.**
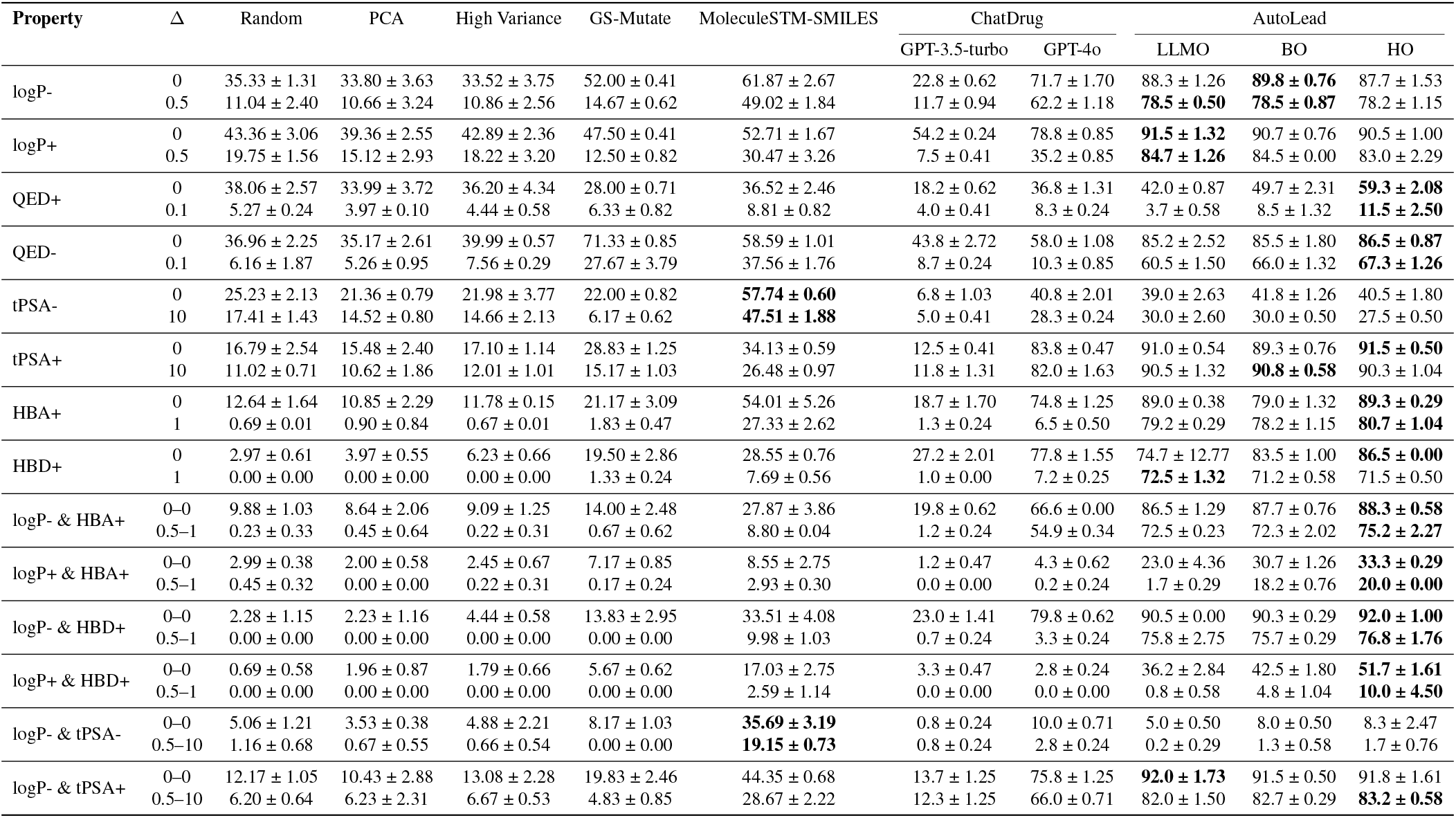
Success rate (%) on 16 ChatDrug-200 single-objective and 12 multi-objective small-molecule editing tasks. Δ denotes the required improvement threshold for each property. Evaluation is the hit ratio of the property change. The best results are marked in **bold**.

In the more demanding 12 multi-objective tasks (in the bottom six rows of Table 1), the superiority of our hybrid approach is further confirmed. The AutoLead framework achieved the best performance on all tasks except for logP& tPSA-, with AutoLead-HO consistently delivering the top performance in the most complex scenarios. While the MoleculeSTM-SMILES baseline is strong in tasks involving tPSA reduction, our method dominates elsewhere. For instance, when simultaneously decreasing logP and increasing hydrogen bond donors (logP-& HBD+) under strict thresholds (Δ = 0.5 − 1), AutoLead-HO’s success rate of **76.8%** is dramatically higher than ChatDrug’s **3.3%**. Likewise, for increasing both logP and HBA+ (Δ = 0.5 − 1), our hybrid method achieves a **20.0%** success rate where the previous state-of-the-art only achieves **0.2%**. For valid rate, please Table S3.

### Performance on the DrugAssist-500 dataset

Results on the DrugAssist-500 dataset (Table S2) further confirm the generalizability and effectiveness of our proposed hybrid method, AutoLead-HO, in interactive multi-property optimization scenarios. The AutoLead framework, including its LLMO and BO variants, demonstrates superior performance by achieving the top success rate across most of the tasks. This outperformance is especially pronounced in the more challenging strict settings. For instance, in the complex multi-property task of decreasing logP while increasing HBA+ with a stringent threshold (Δ = 1), our hybrid approach HO achieves a success rate of 72.0%, a dramatic improvement over the 0.8% from ChatDrug. In another difficult task, increasing the number of hydrogen bond donors (HBD+) with a strict threshold of Δ = 1, our BO variant leads with a 70.7% success rate, again showcasing a substantial leap from the 4.2% achieved by ChatDrug with GPT-4o. For valid rate, please refer Table S4.

### Performance on the LipinskiFix-1000 Dataset

We evaluate AutoLead on the LipinskiFix-1000 dataset, a realistic benchmark that requires methods to restore Lipinski’s Rule of Five compliance while simultaneously improving QED. This task represents a stringent, real-world multi-objective challenge common in early-stage drug discovery.

As summarized in Table 2, our hybrid strategy (AutoLead-HO) demonstrates a significant advantage. It achieves a 28.9% success rate, considerably outperforming previous approaches like MoleculeSTM (13.9%) and ChatDrug (3.87%). In terms of valid rate (Table S5), AutoLead-HO also performs exceptionally well at 93.6%, substantially outperforming methods like MoleculeSTM (49.4%) and ChatDrug (79.3%). The strong performance confirms the framework’s potential for guiding the development of drug-like molecules in realistic lead optimization pipelines.

**Table 2.** Success rate (%) on the LipinskiFix-1000 dataset. The evaluation metrics is the success rate (percentage of molecules that satisfy both Lipinski’s Rule of Five and achieve QED ≥ 0.7). The best results are marked in **bold**.

| Method | Success Rate (%) |
| --- | --- |
| Random | 11.50 ± 3.65 |
| PCA | 11.90 ± 4.04 |
| High Variance | 12.30 ± 4.95 |
| GS-Mutate | 1.50 ± 0.23 |
| MoleculeSTM-SMILES | 13.90 ± 2.55 |
| ChatDrug (GPT-3.5-turbo) | 3.87 ± 0.06 |
| ChatDrug (GPT-4o) | 3.83 ± 0.29 |
| AutoLead-LLMO | 2.50 ± 0.21 |
| AutoLead-BO | 20.30 ± 0.85 |
| AutoLead-HO | **28.33 ± 0.66** |

### Impact of LLM-driven Inverse Mapping

To evaluate the effectiveness of our proposed LLM-driven inverse mapping, we conducted two key analyses.

First, we compared our approach against a conventional deep learning decoder. We replaced the LLM-driven decoder in AutoLead-BO with a Variational Autoencoder (VAE) decoder (30), testing latent space dimensionalities of 16, 32, and 128. As shown in Table S6, when evaluated on representative single- and multi-objective tasks from the ChatDrug-200 dataset (QED+ and logP+ & HBD+), all VAE-based variants underperformed compared to our LLM-driven approach. This result highlights the difficulty of using standard decoders to map from a property-defined latent space back to valid, highquality molecules, validating our novel method.

Second, we sought to confirm that the inverse mapping step itself contributes meaningfully to the optimization process. According to our framework (Algorithm 1), if the LLM-driven decoder fails or generates an invalid SMILES, the system falls back to direct generation via the LLM (*G*_LLM_)- To isolate the contribution of the decoder, we analyzed the performance of molecules generated by each path on the LipinskiFix-1000 benchmark. The results, presented in Table S7, clearly demonstrate that molecules generated via the LLM inverse mapping achieve a significantly higher valid rate and success rate than those produced by the direct LLM generation fallback. This finding confirms that our LLM-driven decoder is a critical component that effectively translates optimized latent vectors into successful molecular candidates, rather than relying on the fallback mechanism.

### Visualization of Successful Lead Optimizations

To further illustrate the practical effectiveness of AutoLead, Figure 2 showcases representative examples of successful lead optimizations. These examples demonstrate AutoLead’s ability to perform complex edits, transforming initial molecules into more favorable candidates by improving key drug-like properties. Notably, we observe significant increases in QED alongside improvements in Lipinski’s Rule of Five compliance. For instance, several transformations show substantial reductions in MW and logP, as well as increases in HBD and HBA counts to fall within desirable ranges. The modifications, such as the strategic introduction of fluorine atoms or subtle structural alterations like ring contractions and functional group removals, are chemically sensible and preserve the core structural motifs of the original molecules. This highlights AutoLead’s capability to generate realistic and relevant chemical transformations aligned with medicinal chemistry principles during early-stage lead optimization. We also provide examples of successful molecule refinements across the ChatDrug-200 and DrugAssist-500 datasets in Figure S2.

**Fig. 2.**
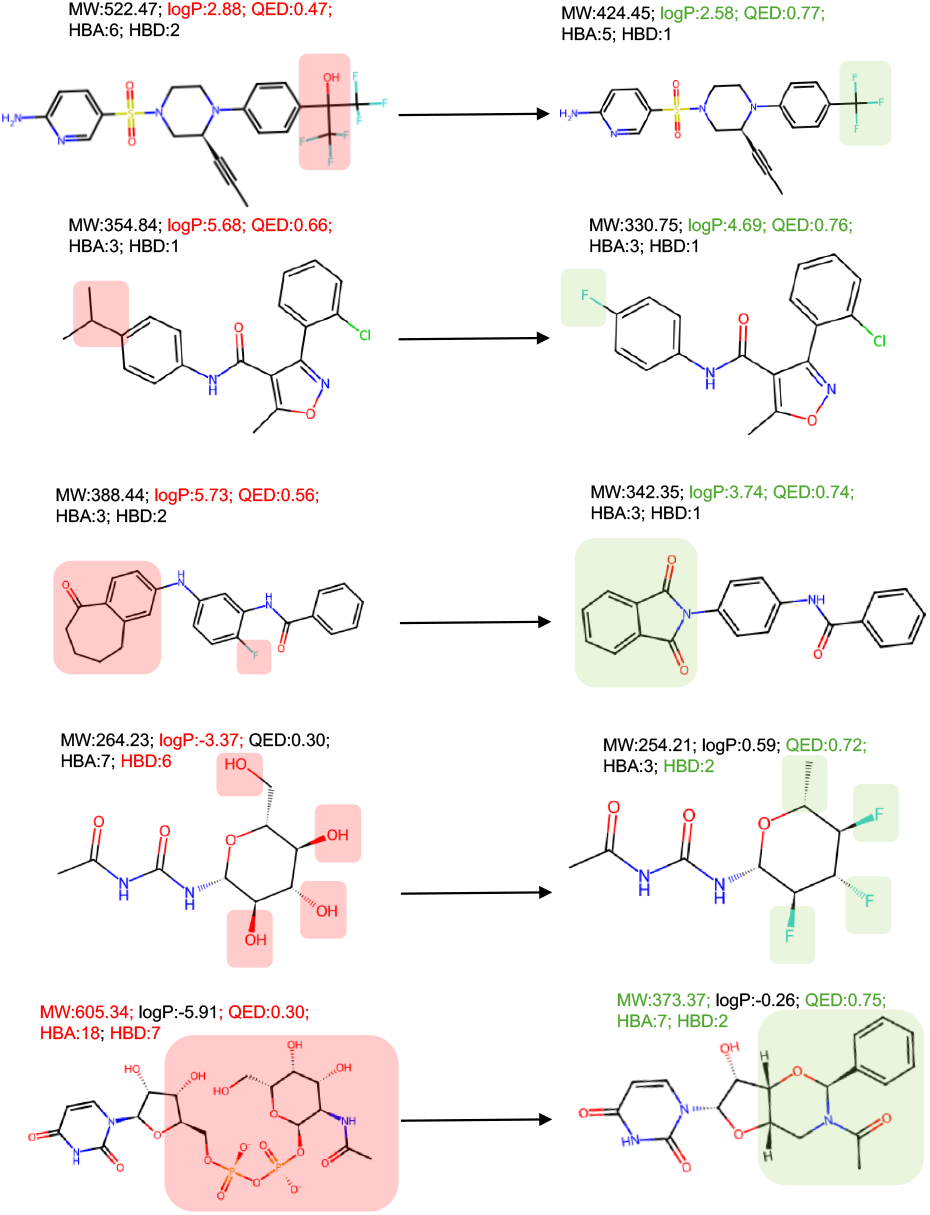
Examples of successful molecule refinement by AutoLead on the LipinskiFix-1000 dataset. For each original (left) and optimized (right) pair, key properties are displayed. Green indicates a favorable change, while red flags the initial property that failed to meet drug-likeness criteria.

### Dynamic Usage of Bayesian Optimization in HO

To understand the contribution of Bayesian Optimization (BO) within our hybrid optimization (HO) strategy, we analyze the proportion of steps where BO was selected during the molecular editing process. The results, summarized in Table S8, reveal that HO dynamically adjusts its reliance on BO based on task difficulty. For simpler property edits like modifying logP+ (Δ = 0), where LLM-based exploration is often sufficient, the BO ratio remains very low (0.17%). In contrast, for tasks requiring more precise, data-driven exploitation, the reliance on BO increases substantially. This is evident in challenging single-objective tasks like QED+ (56.19%) and complex multi-objective cases such as logP+&HBD+ (56.62%). This trend culminates in the LipinskiFix-1000 benchmark, which involves satisfying multiple strict constraints, where BO usage reaches its peak at 64.8%. These findings validate our hybrid design, which effectively balances broad, LLM-driven exploration with focused, BO-driven exploitation according to the complexity of the optimization task.

### Impact of BO Integration and LLM Choice

As shown in Figure S1, integrating Bayesian Optimization (BO) yields substantial performance gains. The hybrid (HO) strategy consistently outperforms the LLM-only (LLMO) approach, regardless of the LLM backbone, while maintaining a high molecule valid rate above 85%.

The analysis also reveals a crucial synergy between the hybrid method and a more advanced LLM. The performance benefit from upgrading the LLM from GPT-3.5-turbo to GPT-4o is magnified under the HO strategy (+2.1%) compared to the modest gain for the LLMO strategy (+0.6%). This indicates that BO’s structured exploration is most effective when paired with a powerful LLM. We hypothesize that such a more capable model serves as a more effective inverse mapping function, 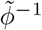, for decoding optimized latent vectors into valid, high-quality molecular structures.

### A Richer Feature Space is Crucial for Optimization Success

We also analyze AutoLead-HO by varying the dimensionality of the BO feature space, comparing a comprehensive 10-dimension (10D) space against smaller 2D and 5D subsets. The 10D space included key descriptors like molecular weight, LogP, TPSA, and hydrogen bond counts. As shown in Table S9, the 10D model performs better on average across the ChatDrug-200 and DrugAssist-500 datasets. The advantage of a richer feature space became most apparent on the more challenging LipinskiFix-1000 benchmark. Here, the 10D model’s success rate of 28.9% represented an order-of-magnitude improvement over the 2D (2.9%) and 5D (2.3%) models. This demonstrates that while a minimal feature set may suffice for simpler tasks, a comprehensive set of molecular descriptors is critical for navigating complex, multi-objective optimization landscapes and achieving robust performance.

## Conclusion

In this work, we introduce AutoLead, a novel framework that integrates the chemical reasoning of LLMs with the principled exploration of multi-objective Bayesian optimization to address realistic lead optimization challenges. Unlike prior approaches that rely on narrow objectives or handcrafted features, AutoLead combines the intuitive generative power of LLMs with the uncertainty-aware search of Gaussian Processes to efficiently explore complex molecular spaces.

Across three diverse benchmarks—ChatDrug-200, DrugAssist-500, and the newly introduced LipinskiFix1000, which simulates a realistic drug-likeness evaluation scenario—AutoLead consistently outperforms existing methods. Further, our hybrid strategy excels on challenging multi-property tasks, surpassing both LLM-only and BO-only baselines. These results underscore the potential of coupling LLMs with black-box optimization for molecular design, paving the way for more robust and scalable solutions in computational drug discovery.

## Supplementary data

Submitted a supplementary material to provide additional results.

## Data availability

The LipinskiFix-1000 data are provided in [figshare](https://figshare.com/articles/dataset/LipinskiFix-1000_dataset/30398953?file=58880725).

## Supplementary Note 1: Prompt Design and Workflow

We employ a novel LLM-in-the-loop multi-objective drug optimization framework that integrates Large Language Models (LLMs) with Bayesian Optimization (BO). The core idea is to use LLMs to generate and refine molecular structures in response to optimization objectives and historical evaluations, while BO guides the search in chemical space.

To facilitate this interaction, we design a set of prompt templates tailored to the distinct stages of the optimization process. Each prompt is carefully crafted to ensure the LLM outputs are chemically valid, optimization-aware, and compatible with downstream evaluation.

Table S1 summarizes the prompt templates used throughout the AutoLead pipeline.

**Table S1.** Prompt templates used across different stages of AutoLead. Each prompt is conditioned on historical data and task objectives, guiding the LLM in molecule generation, transformation, or evaluation.

| Phase | Prompt Template |
| --- | --- |
| Initialization & Molecule Optimization ($G_{LLM}$) | The following are the drug molecules and their properties evaluated in the past: **[History String]**.<br>Optimization objectives: **[Objectives Description]**.<br>Based on the molecule **[Base Molecule SMILES]**, please design an improved new molecule.<br>**IMPORTANT CONSTRAINTS:**<br>You can make the following types of modifications:<br>– ADD functional groups (e.g., -OH, -NH<sub>2</sub>, -COOH, -CH<sub>3</sub>, -F, -Cl)<br>– DELETE existing functional groups or side chains<br>– SUBSTITUTE functional groups (e.g., replace -OH with -NH<sub>2</sub>, -CH<sub>3</sub> with -CF<sub>3</sub>)<br>– Modify ring systems (add/remove substituents on rings)<br>Preserve the core molecular scaffold and main structural framework.<br>Avoid completely changing the molecular backbone or creating entirely different structures.<br>Focus on single-step modifications rather than multiple simultaneous changes.<br>The new molecule should optimize the above objectives while maintaining chemical validity.<br>**PLEASE CAREFULLY CONSIDER THE OPTIMIZATION SUGGESTIONS above when designing the new molecule.**<br>Please only return a valid SMILES string. |
| LLM-driven Inverse Mapping | **[Historical Context]**<br>Based on the above molecular history, please generate a valid SMILES molecule with the following characteristics: **[Feature Description]**.<br>The molecule should be chemically valid and drug-like. Learn from the patterns in the previously generated molecules to create a structurally diverse yet optimized compound.<br>Return only the SMILES string. |

**Table S2.** Success rate (%) on DrugAssist-500 10 small molecule editing. Δ denotes the required improvement threshold for each property. Evaluation is the hit ratio of the property change. The best results are marked in **bold**.

| Property | $\Delta$ | Random | PCA | High Variance | GS-Mutate | MoleculeSTM-SMILES | ChatDrug | | AutoLead | | |
| --- | --- | --- | --- | --- | --- | --- | --- | --- | --- | --- | --- |
|  |  |  |  |  |  |  | GPT-3.5-turbo | GPT-4o | LLMO | BO | HO |
| logP- | 0 | 39.20 ± 3.15 | 42.00 ± 3.40 | 39.60 ± 0.86 | 50.67 ± 0.41 | 53.67 ± 6.03 | 28.2 ± 0.33 | 66.6 ± 0.00 | 86.3 ± 1.29 | 86.6 ± 1.29 | **87.7 ± 0.23** |
| | 0.5 | 25.67 ± 4.06 | 27.20 ± 2.41 | 25.53 ± 1.84 | 13.27 ± 0.66 | 44.20 ± 5.66 | 18.7 ± 0.25 | 54.9 ± 0.34 | 73.5 ± 0.23 | 72.6 ± 0.23 | **75.2 ± 2.27** |
| QED+ | 0 | 44.73 ± 2.21 | 46.47 ± 3.85 | 44.33 ± 3.40 | 30.40 ± 0.91 | 53.80 ± 0.00 | 16.0 ± 0.33 | 28.8 ± 0.49 | 36.1 ± 1.32 | 26.1 ± 16.52 | **62.3 ± 1.75** |
| | 0.1 | 25.20 ± 3.99 | 25.67 ± 2.99 | 22.93 ± 2.47 | 6.00 ± 0.43 | **37.27 ± 6.88** | 6.8 ± 0.28 | 10.1 ± 0.34 | 15.1 ± 0.40 | 10.1 ± 0.50 | 24.3 ± 0.92 |
| HBA+ | 0 | 58.13 ± 1.91 | 58.60 ± 2.55 | 57.80 ± 1.13 | **92.40 ± 1.56** | 63.80 ± 0.00 | 24.1 ± 0.50 | 68.5 ± 1.23 | 86.9 ± 0.50 | 87.3 ± 0.70 | 86.3 ± 0.42 |
| | 1 | 13.13 ± 0.84 | 13.27 ± 0.84 | 12.53 ± 1.67 | 23.93 ± 1.43 | 15.20 ± 8.77 | 0.4 ± 0.16 | 2.1 ± 0.25 | **78.7 ± 0.76** | 77.9 ± 0.61 | 77.1 ± 1.33 |
| HBD+ | 0 | 58.00 ± 2.57 | 58.40 ± 2.47 | 57.67 ± 0.41 | **86.80 ± 1.56** | 68.60 ± 1.98 | 28.6 ± 0.59 | 68.5 ± 0.52 | 84.1 ± 1.03 | 83.2 ± 1.01 | 85.7 ± 0.42 |
| | 1 | 8.00 ± 1.88 | 8.20 ± 1.88 | 8.33 ± 0.57 | 18.60 ± 0.33 | 8.13 ± 1.04 | 0.3 ± 0.09 | 4.2 ± 0.28 | 68.6 ± 0.87 | **70.7 ± 0.60** | 70.2 ± 0.40 |
| logP- & HBA+ | 0-0 | 33.20 ± 1.88 | 35.00 ± 1.56 | 34.40 ± 0.59 | 49.33 ± 0.47 | 38.07 ± 3.49 | 26.8 ± 0.28 | 63.5 ± 1.16 | **84.1 ± 0.95** | 82.0 ± 0.60 | 75.0 ± 0.35 |
| | 0.5-1 | 7.93 ± 0.68 | 9.27 ± 1.09 | 8.47 ± 1.15 | 6.73 ± 0.66 | 11.93 ± 1.32 | 0.2 ± 0.16 | 0.8 ± 0.28 | 70.6 ± 0.53 | 71.0 ± 0.60 | **72.0 ± 0.60** |

**Table S3.**
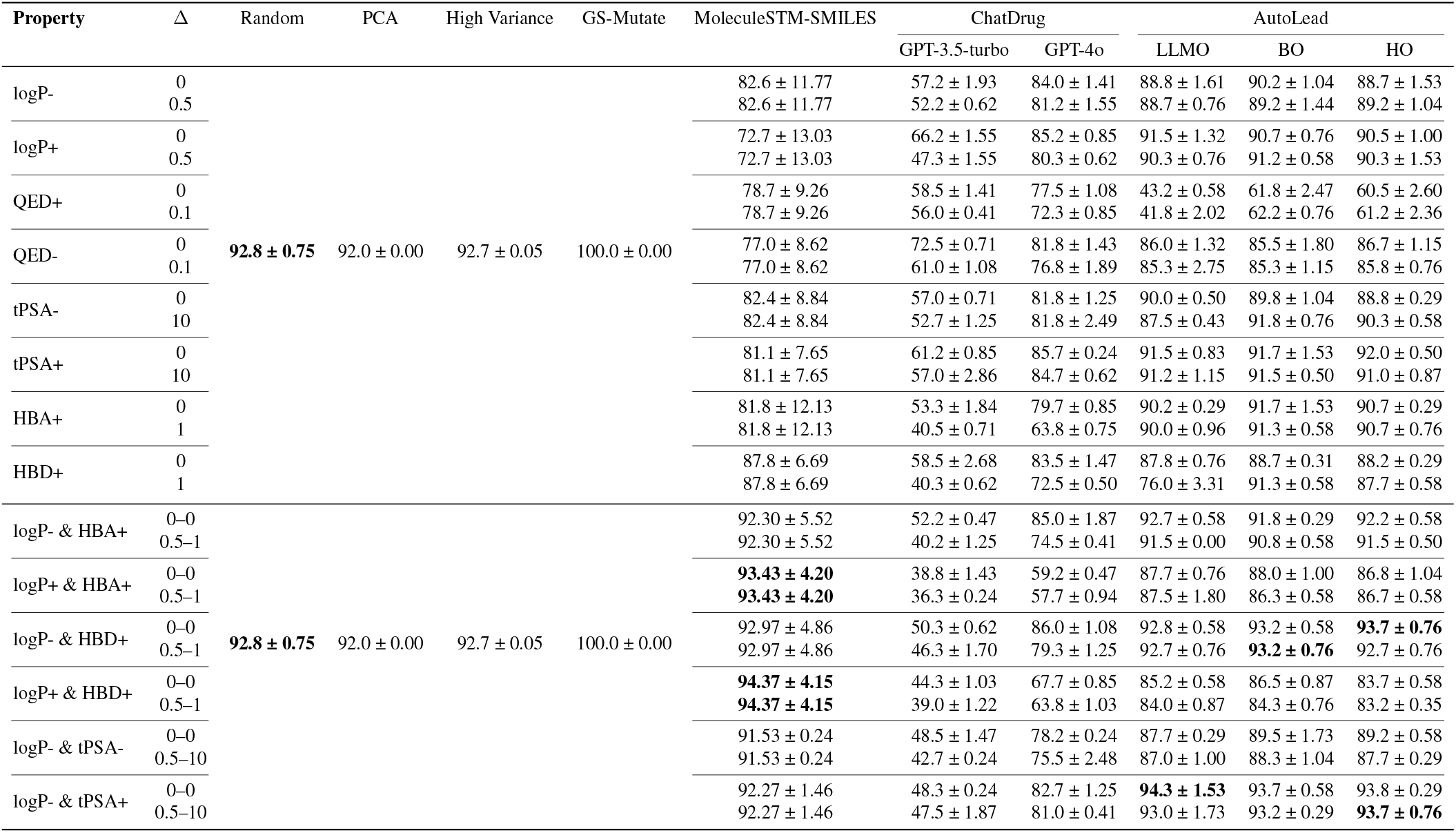
Valid rate (%) on ChatDrug-200 16 single-objective and 12 multi-objective small molecule editing tasks. Evaluation is the hit ratio of the property change. The second-best results are marked in **bold**.

**Table S4.** Valid rate (%) on DrugAssist-500 10 small-molecule editing tasks. Evaluation is the hit ratio of the property change. The second-best results are marked in **bold**.

| Property | $\Delta$ | Random | PCA | High Variance | GS-Mutate | MoleculeSTM-SMILES | ChatDrug | | AutoLead | | |
| --- | --- | --- | --- | --- | --- | --- | --- | --- | --- | --- | --- |
|  |  |  |  |  |  |  | GPT-3.5-turbo | GPT-4o | LLMO | BO | HO |
| logP- | 0 | 70.21 ± 0.38 | 69.57 ± 0.83 | 69.48 ± 0.78 | 100.0 ± 0.00 | 73.87 ± 11.79 | 66.9 ± 1.05 | 82.7 ± 0.68 | 86.9 ± 1.22 | **88.7 ± 0.42** | 88.4 ± 0.35 |
| | 0.5 | | | | | 73.87 ± 11.79 | 64.1 ± 0.94 | 81.8 ± 0.43 | 86.5 ± 1.10 | **87.8 ± 0.69** | 87.3 ± 0.95 |
| QED+ | 0 | | | | | 68.08 ± 4.30 | 69.5 ± 0.74 | **78.1 ± 0.90** | 26.9 ± 16.92 | 63.3 ± 2.04 | 63.3 ± 1.89 |
| | 0.1 | | | | | 68.08 ± 4.30 | 65.1 ± 0.52 | **77.1 ± 0.47** | 46.1 ± 0.58 | 63.6 ± 1.83 | 62.9 ± 1.63 |
| HBA+ | 0 | | | | | 75.61 ± 9.65 | 67.8 ± 0.65 | 80.3 ± 0.41 | 88.8 ± 1.03 | **89.7 ± 0.31** | 88.5 ± 0.81 |
| | 1 | | | | | 75.61 ± 9.65 | 59.3 ± 0.81 | 64.9 ± 0.25 | 88.9 ± 0.12 | **89.3 ± 0.94** | 89.0 ± 1.06 |
| HBD+ | 0 | | | | | 79.52 ± 4.92 | 76.2 ± 0.91 | 79.9 ± 0.68 | 87.5 ± 0.64 | 88.7 ± 0.31 | **89.1 ± 0.12** |
| | 1 | | | | | 79.52 ± 4.92 | 70.7 ± 0.41 | 75.7 ± 1.24 | 88.8 ± 0.40 | 88.7 ± 0.42 | **89.0 ± 0.72** |
| logP- & HBA+ | 0 | | | | | 77.40 ± 5.60 | 63.5 ± 1.15 | 80.8 ± 0.91 | 90.5 ± 0.58 | 91.2 ± 0.20 | **91.4 ± 0.35** |
| | 1 | | | | | 77.40 ± 5.60 | 51.4 ± 0.65 | 69.1 ± 0.90 | 90.7 ± 0.31 | 90.5 ± 0.31 | **90.8 ± 0.72** |

### Initialization of Molecules

Given an initial lead molecule, this stage aims to generate a diverse pool of chemically valid and drug-like candidates using the LLM. The goal is to warm-start the optimization process by producing sufficient data points for fitting the Gaussian Process surrogate model used in subsequent BO steps. The prompt guides the LLM to propose structurally consistent modifications that satisfy domain-specific constraints, such as scaffold preservation and functional group plausibility, without relying on prior optimization history.

### Molecule Optimization

At this stage, the LLM is guided to propose improved molecules based on a given base molecule and a set of optimization objectives (e.g., increase QED or logP). The prompt constrains the model to perform minimal and chemically plausible modifications, such as single-step substitutions or functional group edits, while preserving the core scaffold.

### Feature Vector to SMILES Conversion

During the BO process, molecules are represented as latent feature vectors. This prompt converts such latent feature vectors back into valid SMILES strings by leveraging learned structural patterns from previous generations. The goal is to decode high-quality molecules that exhibit the target features.

#### Valid Rate Analysis Across Benchmarks

Across all evaluated benchmarks: ChatDrug-200 (Table S3), DrugAssist-500 (Table S4), and LipinskiFix-1000 (Table S5), our AutoLead framework consistently demonstrates high chemical validity. The baselines represent different strategies: as a genetic algorithm, GS-Mutate naturally achieves a perfect 100% valid rate, serving as an upper benchmark. In contrast, methods like Random, PCA, High Variance, and MoleculeSTM-SMILES operate by perturbing the latent space of a MegaMolBART autoencoder, which does not guarantee validity upon decoding.

**Table S5.** Valid rate (%) on the LipinskiFix-1000 benchmark.

| Method | Valid Rate (%) |
| --- | --- |
| Random | 48.0 ± 0.17 |
| PCA | 48.3 ± 0.72 |
| High Variance | 48.2 ± 0.37 |
| GS-Mutate | 100.0 ± 0.00 |
| MoleculeSTM-SMILES | 49.4 ± 3.69 |
| ChatDrug (GPT-3.5-turbo) | 70.0 ± 0.59 |
| ChatDrug (GPT-4o) | 79.3 ± 0.32 |
| AutoLead-LLMO | 93.7 ± 0.25 |
| AutoLead-BO | **94.7 ± 0.17** |
| AutoLead-HO | 93.6 ± 1.22 |

**Table S6.** Performance comparison between VAE-based decoders and our proposed LLM-driven inverse mapping on selected tasks from the ChatDrug-200 dataset. We replace the decoder in AutoLead-BO with VAEs of varying latent space dimensions. Both valid rate (%) and success rate (%) are reported.

| Property | $\Delta$ | 16-dim VAE | 32-dim VAE | 128-dim VAE | LLM-driven Inverse |
| --- | --- | --- | --- | --- | --- |
| **Valid Rate (%)** |  |  |  |  |  |
| QED+ | 0 | 40.7 ± 0.29 | 41.8 ± 1.18 | 40.8 ± 1.76 | **61.8 ± 2.47** |
| | 0.1 | 43.0 ± 0.50 | 41.7 ± 1.76 | 43.2 ± 1.53 | **62.2 ± 0.76** |
| logP+ & HBD+ | 0-0 | 86.3 ± 0.29 | 86.2 ± 0.57 | 86.0 ± 1.00 | **86.5 ± 0.87** |
| | 0.5-1 | **87.2 ± 0.29** | 86.8 ± 0.76 | 86.7 ± 0.29 | 84.3 ± 0.76 |
| **Success Rate (%)** |  |  |  |  |  |
| QED+ | 0 | 40.0 ± 0.50 | 40.50 ± 1.32 | 40.17 ± 2.25 | **49.7 ± 2.31** |
| | 0.1 | 3.3 ± 0.58 | 3.2 ± 0.76 | 4.2 ± 1.04 | **8.5 ± 1.32** |
| logP+ & HBD+ | 0-0 | 35.3 ± 1.04 | 39.17 ± 1.89 | 36.5 ± 2.65 | **42.5 ± 1.80** |
| | 0.5-1 | 1.5 ± 0.00 | 1.5 ± 1.32 | 1.3 ± 0.58 | **4.8 ± 1.04** |

**Table S7.** Contribution analysis of the LLM-driven inverse mapping on the LipinskiFix-1000 dataset. We compare the performance of molecules generated via the inverse mapping path against those from the direct LLM generation fallback (*G*_LLM_) for both AutoLead-BO and AutoLead-HO. The results demonstrate that the inverse mapping achieves high valid rates and substantially greater success rates, confirming its effectiveness.

| Method | $G_{LLM}$ | LLM-driven Inverse |
| --- | --- | --- |
| **Valid Rate (%)** |  |  |
| AutoLead-BO | 73.9 ± 0.87 | **78.3 ± 0.85** |
| AutoLead-HO | 74.6 ± 0.72 | **79.0 ± 0.55** |
| **Success Rate (%)** |  |  |
| AutoLead-BO | 6.5 ± 0.35 | **19.5 ± 0.40** |
| AutoLead-HO | 6.6 ± 0.15 | **19.0 ± 0.87** |

**Table S8.** Bayesian-Optimization (BO) selection ratio of the AutoLead-HO across various benchmarks and tasks. The tasks are grouped by dataset and objective type.

| Task / Property | Constraint ($\Delta$) | HO BO-ratio (%) |
| --- | --- | --- |
| **ChatDrug-200 – Single-Objective Tasks** |  |  |
| logP- | 0 | 3.17 |
|  | 0.5 | 11.16 |
| logP+ | 0 | 0.17 |
|  | 0.5 | 2.42 |
| QED+ | 0 | 33.81 |
|  | 0.1 | 56.19 |
| QED- | 0 | 8.67 |
|  | 0.1 | 35.01 |
| tPSA- | 0 | 35.16 |
|  | 10 | 43.46 |
| tPSA+ | 0 | 0.00 |
|  | 10 | 0.00 |
| HBA+ | 0 | 1.63 |
|  | 1 | 11.50 |
| HBD+ | 0 | 1.58 |
|  | 1 | 12.31 |
| **ChatDrug-200 – Multi-Objective Tasks** |  |  |
| logP- & HBA+ | 0–0 | 2.54 |
|  | 0.5–1 | 12.86 |
| logP+ & HBA+ | 0–0 | 30.33 |
|  | 0.5–1 | 37.79 |
| logP- & HBD+ | 0–0 | 1.46 |
|  | 0.5–1 | 14.03 |
| logP+ & HBD+ | 0–0 | 42.10 |
|  | 0.5–1 | 56.62 |
| logP- & tPSA- | 0–0 | 59.86 |
|  | 0.5–10 | 61.78 |
| logP- & tPSA+ | 0–0 | 1.81 |
|  | 0.5–10 | 4.56 |
| **DrugAssist-500 Tasks** |  |  |
| logP- | 0 | 4.80 |
|  | 0.5 | 14.29 |
| QED+ | 0 | 30.51 |
|  | 0.1 | 52.88 |
| HBA+ | 0 | 2.30 |
|  | 1 | 13.59 |
| HBD+ | 0 | 3.50 |
|  | 1 | 13.93 |
| logP- & HBA+ | 0 | 17.34 |
|  | 1 | 19.20 |
| **LipinskiFix-1000 Dataset** |  |  |
| Overall task |  | 64.80 |

**Table S9.**
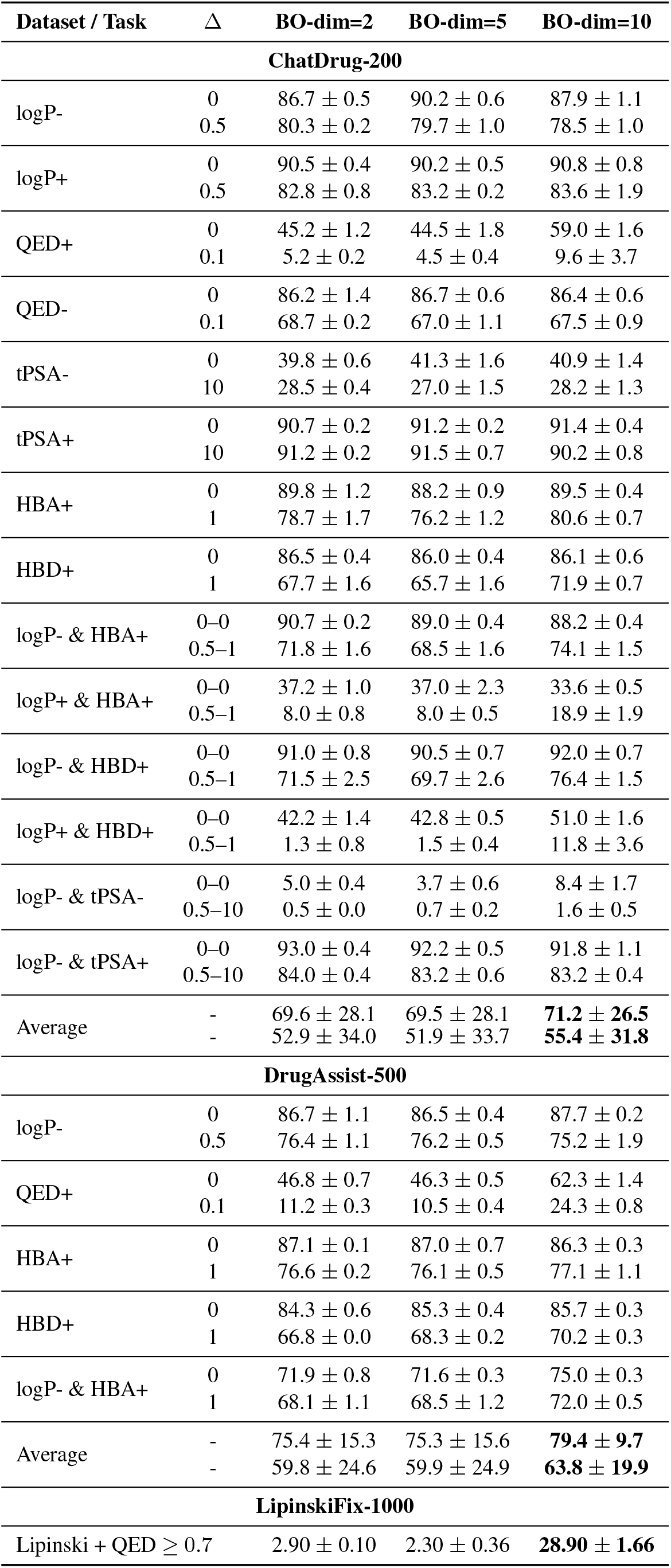
Success rate (%) of AutoLead-HO with different BO latent dimensions (2, 5, 10). Δ denotes the required improvement threshold for each property.

| Dataset / Task | $\Delta$ | BO-dim=2 | BO-dim=5 | BO-dim=10 |
| --- | --- | --- | --- | --- |
| **ChatDrug-200** |  |  |  |  |
| logP- | 0 | 86.7 ± 0.5 | 90.2 ± 0.6 | 87.9 ± 1.1 |
| | 0.5 | 80.3 ± 0.2 | 79.7 ± 1.0 | 78.5 ± 1.0 |
| logP+ | 0 | 90.5 ± 0.4 | 90.2 ± 0.5 | 90.8 ± 0.8 |
| | 0.5 | 82.8 ± 0.8 | 83.2 ± 0.2 | 83.6 ± 1.9 |
| QED+ | 0 | 45.2 ± 1.2 | 44.5 ± 1.8 | 59.0 ± 1.6 |
| | 0.1 | 5.2 ± 0.2 | 4.5 ± 0.4 | 9.6 ± 3.7 |
| QED- | 0 | 86.2 ± 1.4 | 86.7 ± 0.6 | 86.4 ± 0.6 |
| | 0.1 | 68.7 ± 0.2 | 67.0 ± 1.1 | 67.5 ± 0.9 |
| tPSA- | 0 | 39.8 ± 0.6 | 41.3 ± 1.6 | 40.9 ± 1.4 |
| | 10 | 28.5 ± 0.4 | 27.0 ± 1.5 | 28.2 ± 1.3 |
| tPSA+ | 0 | 90.7 ± 0.2 | 91.2 ± 0.2 | 91.4 ± 0.4 |
| | 10 | 91.2 ± 0.2 | 91.5 ± 0.7 | 90.2 ± 0.8 |
| HBA+ | 0 | 89.8 ± 1.2 | 88.2 ± 0.9 | 89.5 ± 0.4 |
| | 1 | 78.7 ± 1.7 | 76.2 ± 1.2 | 80.6 ± 0.7 |
| HBD+ | 0 | 86.5 ± 0.4 | 86.0 ± 0.4 | 86.1 ± 0.6 |
| | 1 | 67.7 ± 1.6 | 65.7 ± 1.6 | 71.9 ± 0.7 |
| logP- & HBA+ | 0-0 | 90.7 ± 0.2 | 89.0 ± 0.4 | 88.2 ± 0.4 |
| | 0.5-1 | 71.8 ± 1.6 | 68.5 ± 1.6 | 74.1 ± 1.5 |
| logP+ & HBA+ | 0-0 | 37.2 ± 1.0 | 37.0 ± 2.3 | 33.6 ± 0.5 |
| | 0.5-1 | 8.0 ± 0.8 | 8.0 ± 0.5 | 18.9 ± 1.9 |
| logP- & HBD+ | 0-0 | 91.0 ± 0.8 | 90.5 ± 0.7 | 92.0 ± 0.7 |
| | 0.5-1 | 71.5 ± 2.5 | 69.7 ± 2.6 | 76.4 ± 1.5 |
| logP+ & HBD+ | 0-0 | 42.2 ± 1.4 | 42.8 ± 0.5 | 51.0 ± 1.6 |
| | 0.5-1 | 1.3 ± 0.8 | 1.5 ± 0.4 | 11.8 ± 3.6 |
| logP- & tPSA- | 0-0 | 5.0 ± 0.4 | 3.7 ± 0.6 | 8.4 ± 1.7 |
| | 0.5-10 | 0.5 ± 0.0 | 0.7 ± 0.2 | 1.6 ± 0.5 |
| logP- & tPSA+ | 0-0 | 93.0 ± 0.4 | 92.2 ± 0.5 | 91.8 ± 1.1 |
| | 0.5-10 | 84.0 ± 0.4 | 83.2 ± 0.6 | 83.2 ± 0.4 |
| Average | - | 69.6 ± 28.1 | 69.5 ± 28.1 | **71.2 ± 26.5** |
| | - | 52.9 ± 34.0 | 51.9 ± 33.7 | **55.4 ± 31.8** |
| **DrugAssist-500** |  |  |  |  |
| logP- | 0 | 86.7 ± 1.1 | 86.5 ± 0.4 | 87.7 ± 0.2 |
| | 0.5 | 76.4 ± 1.1 | 76.2 ± 0.5 | 75.2 ± 1.9 |
| QED+ | 0 | 46.8 ± 0.7 | 46.3 ± 0.5 | 62.3 ± 1.4 |
| | 0.1 | 11.2 ± 0.3 | 10.5 ± 0.4 | 24.3 ± 0.8 |
| HBA+ | 0 | 87.1 ± 0.1 | 87.0 ± 0.7 | 86.3 ± 0.3 |
| | 1 | 76.6 ± 0.2 | 76.1 ± 0.5 | 77.1 ± 1.1 |
| HBD+ | 0 | 84.3 ± 0.6 | 85.3 ± 0.4 | 85.7 ± 0.3 |
| | 1 | 66.8 ± 0.0 | 68.3 ± 0.2 | 70.2 ± 0.3 |
| logP- & HBA+ | 0 | 71.9 ± 0.8 | 71.6 ± 0.3 | 75.0 ± 0.3 |
| | 1 | 68.1 ± 1.1 | 68.5 ± 1.2 | 72.0 ± 0.5 |
| Average | - | 75.4 ± 15.3 | 75.3 ± 15.6 | **79.4 ± 9.7** |
| | - | 59.8 ± 24.6 | 59.9 ± 24.9 | **63.8 ± 19.9** |
| **LipinskiFix-1000** |  |  |  |  |
| Lipinski + QED ≥ 0.7 | | 2.90 ± 0.10 | 2.30 ± 0.36 | **28.90 ± 1.66** |

Among the generative models, our AutoLead variants consistently achieve higher valid rates than the LLM-based ChatDrug baselines. On the ChatDrug-200 tasks, AutoLead maintains valid rates often exceeding 90%. This performance is substantially higher than that of ChatDrug (GPT-4o), whose rates are often below 85%, and particularly ChatDrug (GPT-3.5-turbo), which frequently scores below 55%. This trend continues on the DrugAssist-500 benchmark, where AutoLead’s validity remains robust; for example, the HO variant reaches 91.4% for the logP- & HBA+ task. On the stricter LipinskiFix-1000 benchmark, AutoLead’s strong generalization is evident, with the BO variant achieving a 94.7% valid rate, substantially higher than ChatDrug’s 79.3%. This consistent high performance in generating chemically valid molecules underscores the reliability of the AutoLead framework for practical drug design applications.

**Fig. S1.**
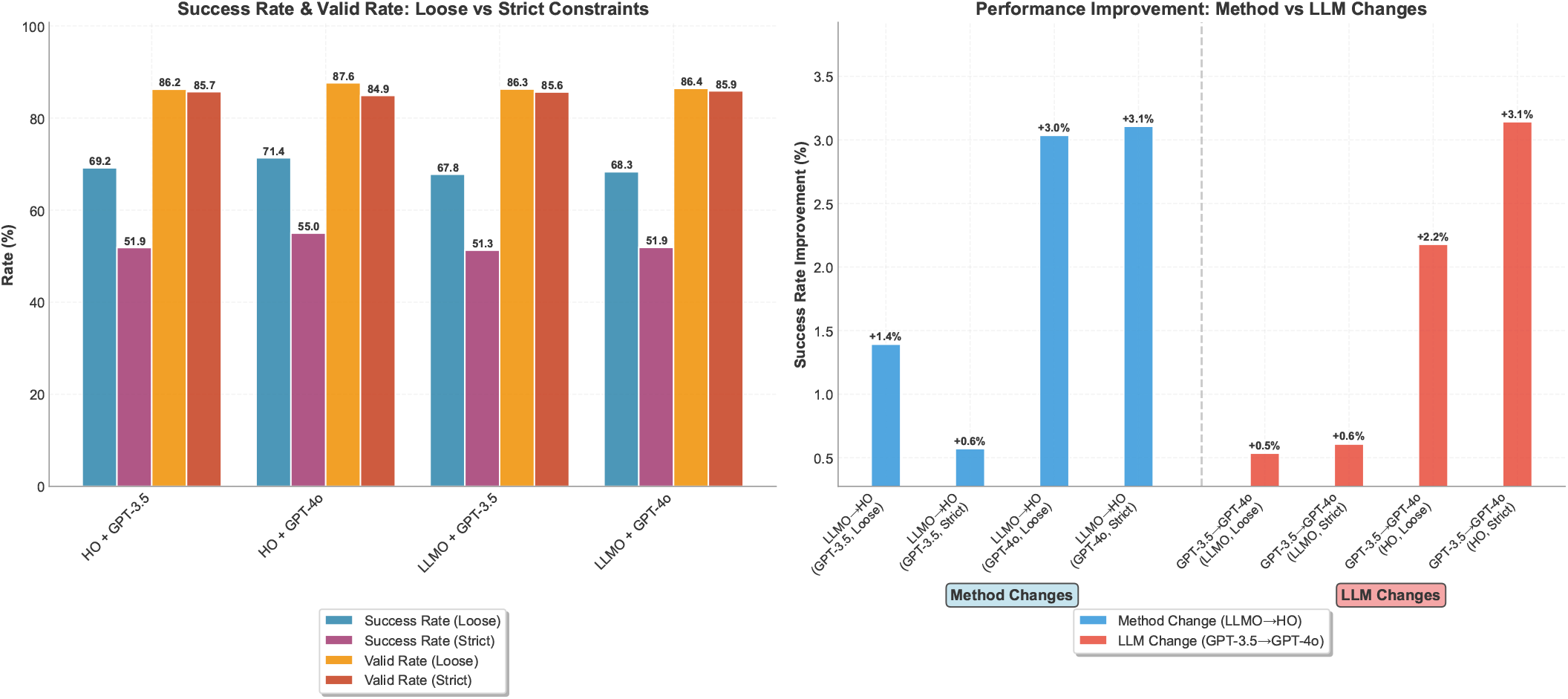
Performance comparison on the ChatDrug-200 dataset. **Left:** Success rates and valid rates under both loose and strict constraint settings for different variants of HO and LLMO. **Right:** Success rate improvements attributable to method changes (LLMO → HO) versus LLM backbone changes (GPT-3.5 → GPT-4o). These results highlight the relative contribution of each factor to overall performance gains.

**Fig. S2.**
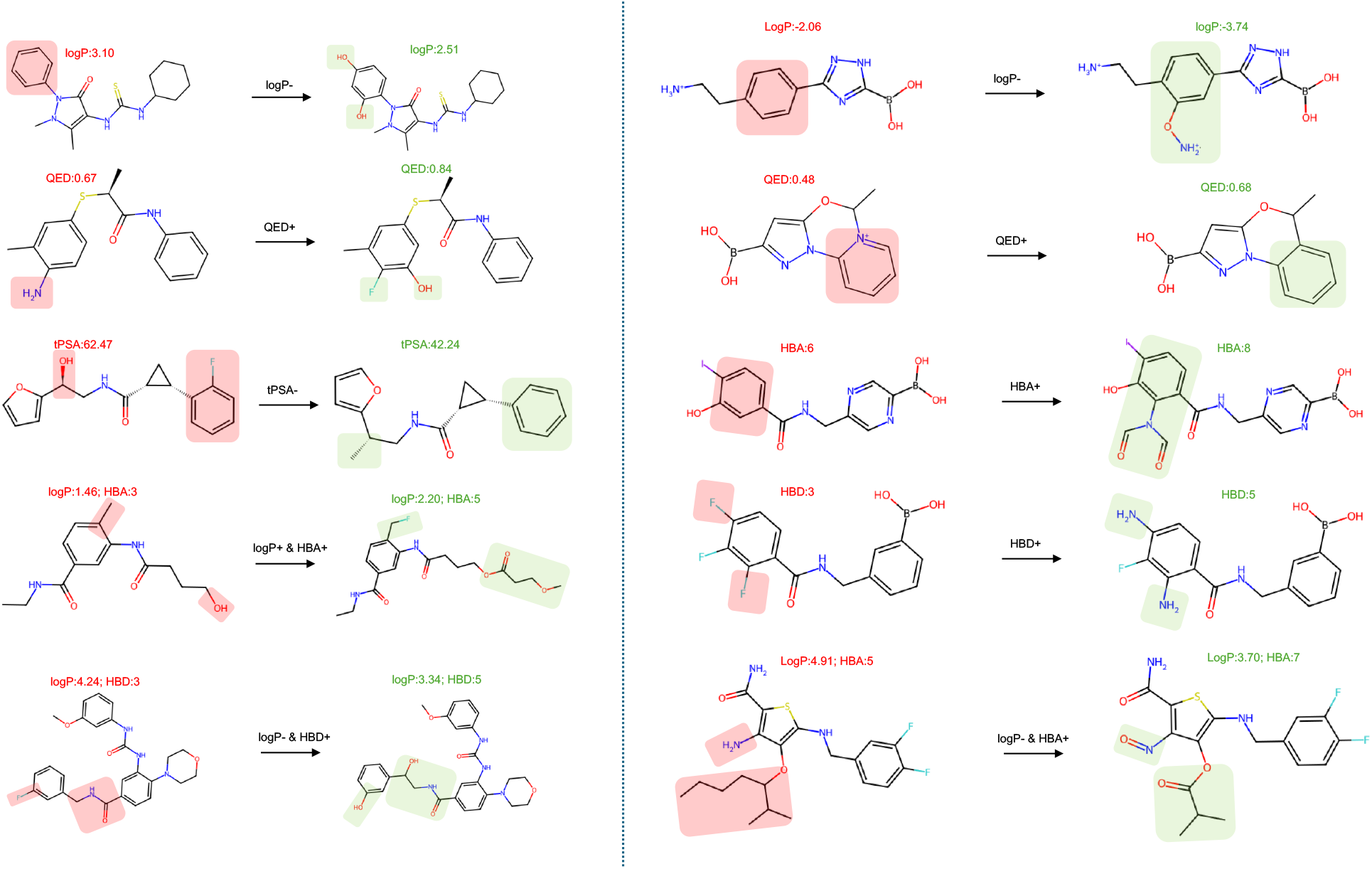
Examples of molecule refinements generated by AutoLead across the ChatDrug-200 and DrugAssist-500 datasets. Each pair shows the original molecule (left) and the optimized molecule (right) with corresponding key property values. Red highlights indicate properties or structural regions in the initial molecule that violated optimization criteria, while green highlights show improvements or favorable modifications introduced by AutoLead.

